# A Novel Strategy for Dynamic Modelling of Genome-Scale Interaction Networks

**DOI:** 10.1101/2022.05.20.491854

**Authors:** Pooya Borzou, Jafar Ghaisari, Iman Izadi, Yasin Eshraghi, Yousof Gheisari

**Author notes:** Corresponding author;, Jafar Ghaisari.

## Abstract

**Background:** Modern medicine is equipped with huge amounts of big biological datasets and a wide range of computational methods to understand the molecular events underlying complex disorders. The recent availability of omics data allows a holistic view towards the interactions of various biomolecule types. However, the constructed maps are static, ignoring the dynamicity of molecular processes. On the other hand, the dynamic models of biological systems are commonly generated in small scales. Hence, the construction of large scale dynamic models that can quantitatively predict the time-course cellular behaviors is a big challenge. This study was aimed at developing a pipeline for automatic construction of such models from time-course experimental data.

**Results:** Information of interactions between input genes is retrieved from SIGNORE 2.0 database and an interaction network is constructed which then is translated to biochemistry language and converted to a biochemical reactions network. In the next step, a large-scale ODE system is constructed by generating the ODE equivalent of each biochemical reaction. To estimate the kinetics parameters of the ODE model, a novel large-scale parameter approximation method is proposed. This method gives an estimation of system parameters by fitting model outputs to time-course experimental measurements. The total pipeline is provided as a MATLAB toolbox called SPADAN, standing for Systematic Protein Association Dynamic ANalyzer. Using multilayer time-series experimental data, the performance of the pipeline was checked by modeling 4379 regulatory interactions between 768 molecules in colon cancer cells exposed to chemotherapy agents.

**Conclusion:** Starting from time-series experimental data, SPADAN automatically constructs map of interactions, generates an ODE system, and performs a parameter approximation procedure. It constructs genome-scale dynamic models, filling the gap between large-scale static and small-scale dynamic modeling strategies. This simulation approach allows for holistic quantitative predictions which is critical for the simulation of therapeutic interventions in precision medicine.

## Introduction

Although in the past decades, medicine has experienced great success in different areas such as infectious diseases, surgical procedures, imaging techniques, and diagnostic tests, efficient management of most non-communicable disorders (NCDs) is yet an unmet goal. The recent availability of huge amount of data has led to the understanding that most disorders are not caused by malperformance of a few genes or proteins, but alteration of the interactions between a large number of biomolecules [1]. The emergence of systems biology has raised hopes to approach the complexity of the underlying mechanisms of chronic diseases and pave the way for more satisfying therapeutic strategies.

In the top-down approach of systems biology, graph theory is employed to analyze the map of interactions between large numbers of biomolecules[2]. However, these networks are static and provide only a snapshot of the system, ignoring the time-dependency nature of biomedical processes. On the other hand, dynamic modeling strategies employed in the bottom-up approach of systems biology allow the generation of quantitative and predictive models which incorporate the dynamism of such processes[3], [4]. Although such models are valuable tools to analyze and forecast the functions of biological systems, they are generally constructed in small scales. Indeed, the generation of predictive models of diseases which are both dynamic and holistic is yet a major challenge.

The generation of large-scale dynamic models of metabolic systems has been attempted by some investigators; Smallbone et al developed a dynamic model of yeast metabolic machinery consisting of 820 metabolites and 956 metabolic reactions[5]. The model structure was constructed by appending some previous dynamic models and the kinetic parameters were harvested from the Biomodels database repository [5]. To overcome the problems related to the large-scale of genome-wide metabolic networks, Smith et al presented a python package, named DMPy, which uses metabolic networks (containing the details of biochemical reactions) as input and automatically converts it to a large set of differential equations. Then, the parameter values are collected from different databases to construct the dynamic model [6]. In addition to ODEs, constraint-based modeling which imposes known biological constraints to limit the solution space [7], has also been employed to model large-scale metabolic networks [8], [9].

The partial adequacy of knowledge on the rate of enzymatic reactions in metabolic networks has made the generation of genome-wide dynamic models feasible. However, to the best of our knowledge, such models are not yet developed for protein-protein interaction (PPI) networks. Not only the lack of sufficient experimental evidence on the rate of reactions, but also the unavailability of large-scale biochemical information of the reactions has hindered the conversion of big PPI networks to dynamic models.

In this paper, we introduce a novel strategy to automatically convert PPI networks to genome-wide dynamic models. In this pipeline, for a given list of proteins, the PPI network is constructed and then the details of reactions are considered in order to translate the network data to the biochemical language. This biochemical network is then automatically converted to an ODE system. In the next step of the pipeline, a parameter estimation algorithm based on a large-scale and distributed approach is proposed to provide system parameters according to high throughput time-series experimental data. This strategy, which is called SPADAN, allows a holistic insight into the dynamism of protein interactions and provides quantitative predictions of system behavior. We have assessed the applicability of this approach to model the interactions of proteins in colorectal cancer and to predict response to specific chemotherapy agents.

## Methods

### Data acquisition

A proteomics dataset (PXD007740) pertinent to the time-course exploration of proteomics and phosphoproteomics of colorectal cancer cells produced by Anna Ressa et al [10]. was retrieved from the ProteomeXchange database^1^. The analyzed RNAseq data of these cells were also obtained as a supplementary file of this report [10].

### Proteomics and Phosphoproteomics data analysis

Raw MS data were analyzed with MaxQuant version (1.6.8.0) integrated with Andromeda search engine against human-reviewed proteome from UniProt FASTA database. Trypsin was configured as a specific enzyme with a maximum of two missed cleavages. For proteomics data, cysteine carbamidomethylation was considered as fixed modification and methionine oxidation and N-terminal acetylation as variable modifications. For phosphopeptides, cysteine carbamidomethylation and phospho (STY) were selected. Proteins were quantified based on unique+razor peptides and two minimum ratio counts. A significance threshold of 0.1 was considered both for peptide spectrum match and protein false discovery rates. The “match between runs” was enabled for all analyses. All quantified peptides and phosphopeptides were filtered for reverse, contaminant, and only identified by sites. Also, phosphosites were retained if they were below a localization probability rate of 75%. To estimate the absolute abundance of proteins “proteomic ruler” plugin of Perseus was employed.

### Hardware description

SPADAN was written and developed in MATLAB 2015b. The SPADAN computation procedure is performed by a PC with16GB RAM and Intel(R) Core(™) i3-6100 CPU.

## Results

This study is aimed at developing a strategy to generate dynamic insights of interactions at the proteome scale. Starting from time-series experimental data, the developed algorithm generates a PPI network and then translates the interactions into a biochemical language. This biochemical network is then converted to a series of ODEs and parameters are estimated using a novel large-scale and distributed parameter estimation technique. The algorithm performs all of these steps automatically. This pipeline is schematically depicted in Fig. 1 and the steps are described below.

**Fig. 1.**
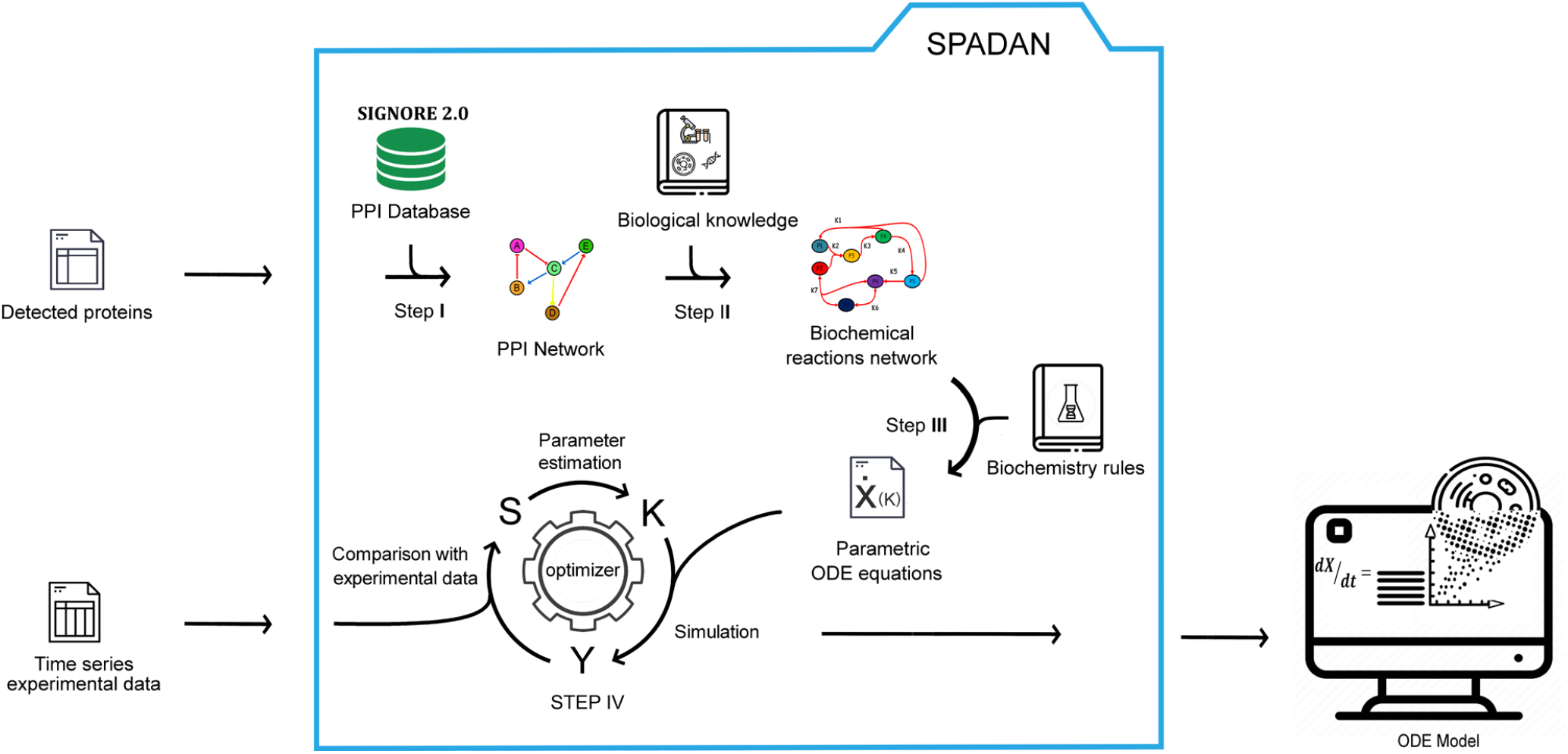
SPADAN modeling procedure pipeline. SPADAN toolbox uses high throughput time-series experimental data from cells to make an ODE model from them. From step I to step III, SPADAN only uses IDs of gene elements detected in the considered cell. In step IV, the values of time-series concentrations are used to estimate unknown parameters of the ODE model.

### Step I: Harvesting network interactions

After getting the list of proteins, SPADAN harvests interactions from the SIGNORE database (ppr83), which includes experimentally validated protein interactions. For the scale of feasibility, we focused on common interaction types including phosphorylation, dephosphorylation, ubiquitination, binding, and transcriptional regulation. An advantage of the SIGNORE database is that the effect of interactions on involved proteins (activation, inhibition, upregulation, or downregulation) is known which is essential for the construction of biochemical networks. The PPI network from SPADAN gives a holistic static view of the interactions between given proteins.

### Step II: Converting PPI network to biochemical reaction network

The construction of PPI networks is a way to schematically illustrate the interactions between proteins. This depiction mode is concise and mainly focused on outcomes rather than processes. For instance, phosphorylation of protein A by protein B is simply shown with a single edge, ignoring different underlying molecular processes. However, in terms of biochemical processes, this interaction is a set of reactions each with a different parameter. Similarly, other molecular interactions can be considered as a series of biochemical reactions (Table 2). This translation from network language to biochemical reactions is an essential step in the construction of the model structure. In order to include transcriptional regulations, the participating genes are considered at DNA and RNA levels in addition to the protein level. For instance, in transcriptional upregulation, regulatory protein A binds the DNA of gene B and activates the transcription of B RNA which then is translated to protein B. For genes without transcriptional regulation edges, a basal level of protein production is assumed. In addition, a degradation rate is assumed for all reactants and products. An example of converting a small PPI network to biochemical reactions network is shown in Fig. 2.

**Fig. 2.**
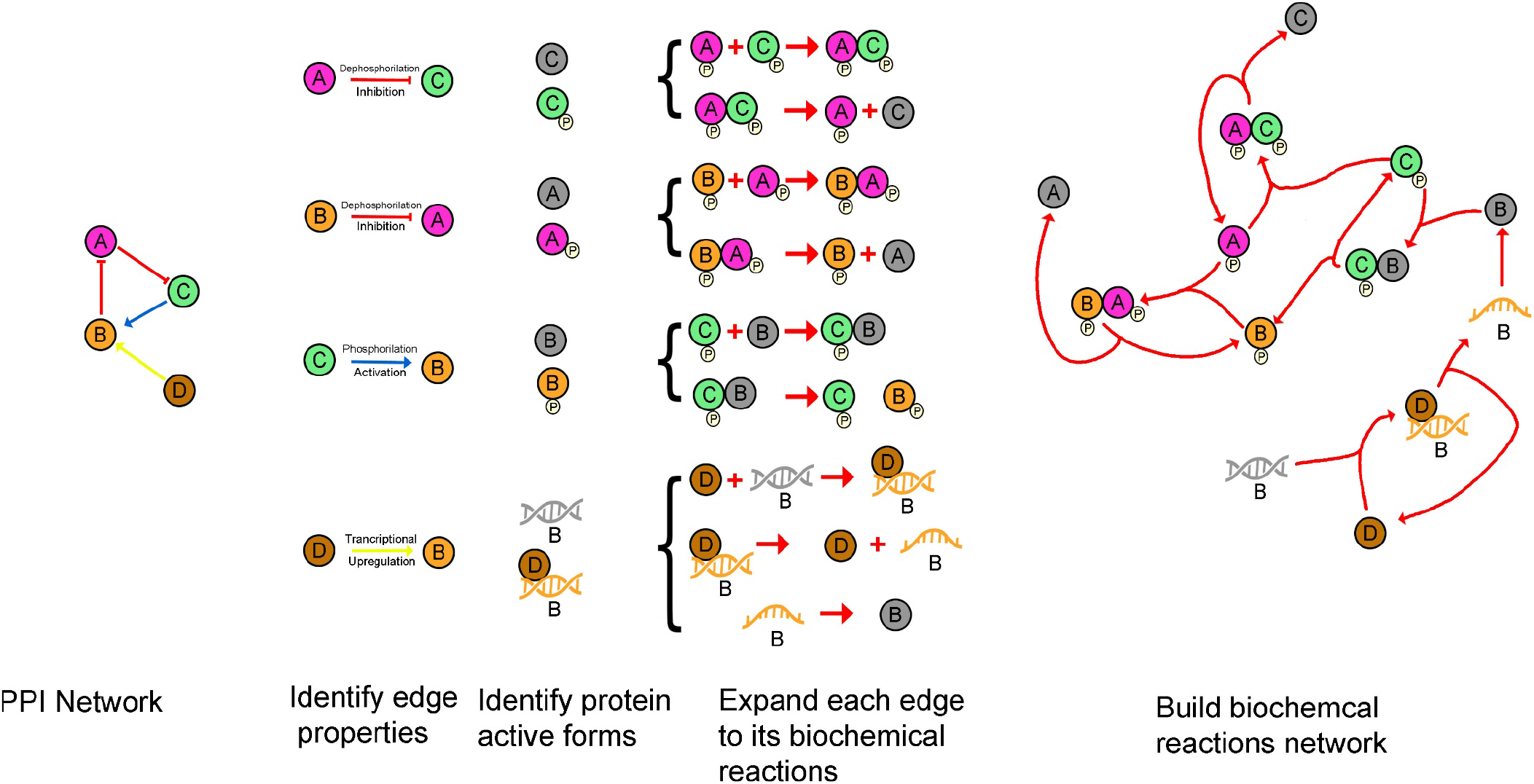
An example from converting PPI network to biochemical reactions network. The shown procedure is performed by SPADAN automatically to use the biochemical reactions network in further modeling steps.

Using the data harvested from SIGNORE, SPADAN generates an “active mode table” showing different forms of proteins in terms of PTM modifications and indicates which PTM mode is the active state of each protein (Table 1). We acknowledge that this is a simplification of the real world in which PTM modifications do not necessarily result in on and off switch-like behaviors but may result in partial augmentation or inhibition of basal activities and also the active state is not restricted to one of the PTM modes.

**Table 1.** Active mode table. In the modeling procedure performed by SPADAN, for each protein, only one of three forms shown in the table is consumed as active and the other two are consumed as inactive forms. The table is automatically made by scanning the types and effects of PPI network edges.

| ID | protein | phosphorilated protein | ubiquitinated protein |
| --- | --- | --- | --- |
| proteini 50 | 0 | 1 | 0 |
| proteini 51 | 1 | 0 | 0 |
| proteini 52 | 1 | 0 | 0 |
| proteini 53 | 0 | 1 | 0 |
| proteini 54 | 0 | 1 | 0 |
| proteini 55 | 0 | 1 | 0 |
| proteini 56 | 1 | 0 | 0 |
| proteini 57 | 1 | 0 | 0 |
| proteini 58 | 1 | 0 | 0 |
| proteini 59 | 0 | 1 | 1 |
| proteini 60 | 0 | 1 | 0 |
| proteini 61 | 1 | 0 | 0 |
| proteini 62 | 0 | 1 | 0 |
| proteini 63 | 1 | 0 | 0 |
| proteini 64 | 1 | 0 | 0 |
| proteini 65 | 0 | 1 | 0 |

**Table 2.**
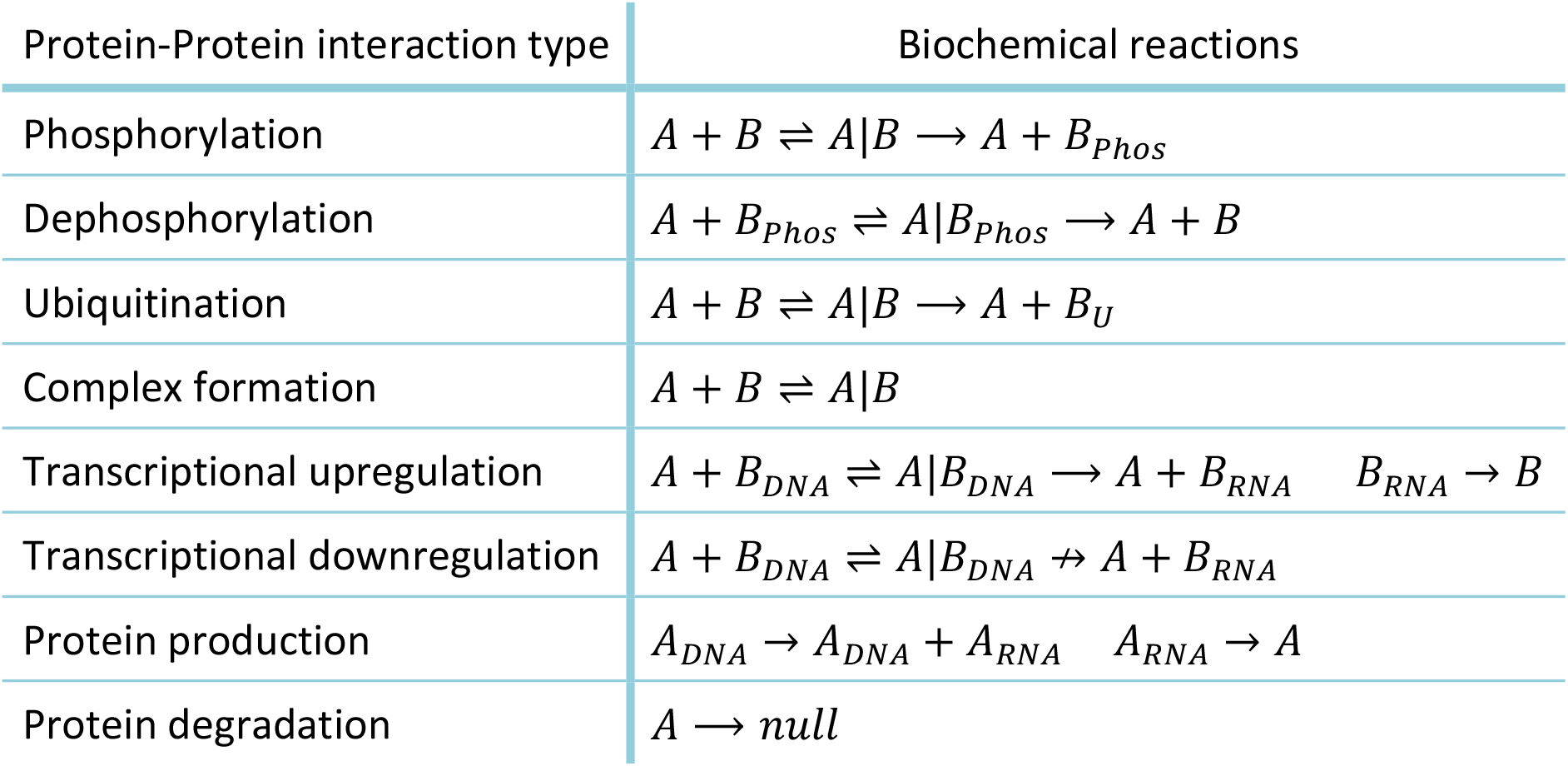
Simplified Biochemical reactions that each type of Protein-Protein interaction includes.

An advantage of the developed algorithm is that it recognizes that each protein is present in different fractions at a given time point in terms of PTM modifications, bound to DNA or bound to other proteins. Indeed, the measured concentration of a given protein at a time is the sum of concentrations of these fractions. Each protein form is considered as an element and receives a unique ID in the algorithm. Furthermore, each element may participate in different reactions and there are overlaps between reactants and products of different reactions. Hence, the biochemical reactions are inter-connected and construct a biochemical network. In order to organize the data of metabolic reaction network for downstream computational processes, a “3D reactions matrix” is generated with several 2D pages. Each page represents a biochemical reaction with two columns for reactants and products. The matrix is shown in Fig. 3.

**Fig. 3.**
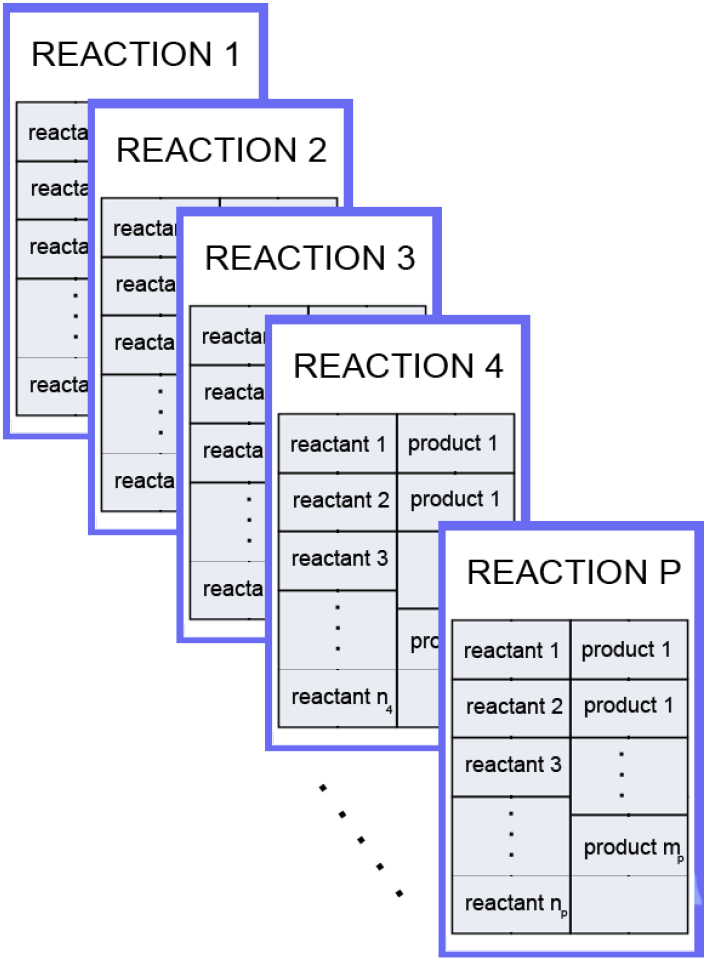
The 3D matrix “reactions” is used by SPADAN to store biochemical reactions network. Each page of the matrix represents one reaction in the biochemical reactions network that includes IDs of reactants and products in the first and second columns respectively. These two columns do not necessarily have same size.

### Step III: Converting biochemical reactions network to a large scale ODE system

The biochemical reactions network provides deep insight on what is happening between interacting molecules. However, it does not explain the dynamic behavior of the system. In order to fuse dynamic insights, a large-scale ODE system is constructed by generating the ODE equivalent of each biochemical reaction automatically. Although SPADAN is able to employ Michaels-Menten or Hill kinetics, all reactions in the colorectal cancer biochemical network are modeled based on mass action law in order to avoid pre-assumptions of those models. Consequently, the changing speed of molecules’ concentration in each reaction is written as a function of the concentrations of substrates and the kinetic parameter of the reaction. Since each molecule can take part in several reactions, the rate of the concentration change for it is the algebraic sum of the concentration changing rates in those reactions. PPI networks are generally constructed to provide a holistic description of the underlying events of complex biomedical phenomena, commonly composed of large numbers of interactions. The size of the model even expands when PPI networks are translated to biochemical networks. Thus, the ODE system constructed based on these networks has an unusually large scale. Therefore, applying novel strategies to decrease computational costs is of critical importance.

Since the ODE solver used in this paper employs the Euler method which is explained in (1), the solver calls function *F*in each time step of integration.

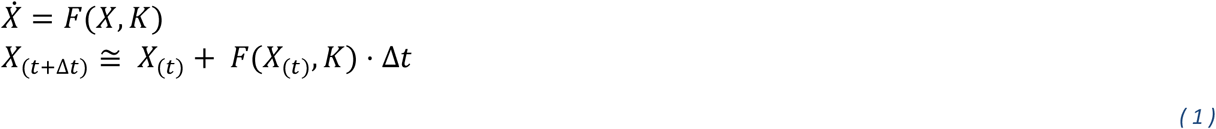

where *X* and *K* are the matrices of state variables and model parameters. The function *F*is the matrix of parametric state equtions of the model which depends on *K* and *X*.

On the other hand, completing the integration requires thousands of time steps. Thus, decreasing the computation cost of the function *F*has a considerable effect on the progression speed of model simulation and consequently in parameter estimation. In order to lower the computation cost of calculating *F*, the SPADAN converts the nonlinear ODE system to matrix multiplications as described further.

Since mass-action law is employed to analyze reaction speeds, the rate of reaction *i* which is shown with *v*_*i*_ can be shown as follow:

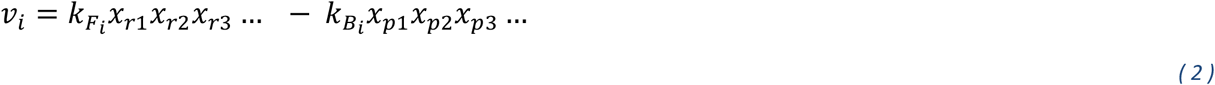

where 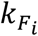 and 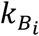 are forward and backward reaction rate constants, *x*_*r1*_, *x*_*r2*_, *x*_*r3*_,… represent reactants of the reaction and *x*_*p1*_, *x*_*p2*_, *x*_*p3*_,… represent the products.

The derivative of each state variable is calculated by

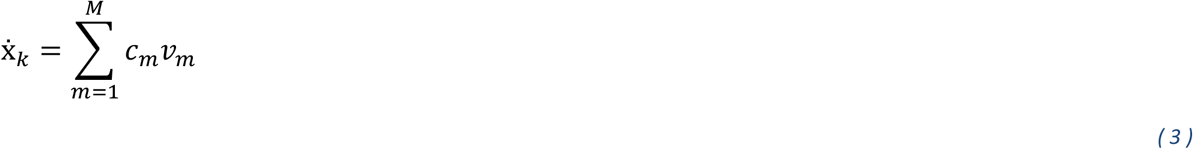

where *M* is the total number of reactions, *v*_*m*_ is the rate of the reaction m and *c*_*m*_ is a variable with 3 possible values of 0, 1, or -1. The variable *c*_*m*_ is evaluated based on the role of the state variable *x*_*k*_ in the reaction *m*.

The nonlinear structure of the equations in (2) and (3) shows that it is not possible to write the ODE system in the form of a linear ODE system as follows:

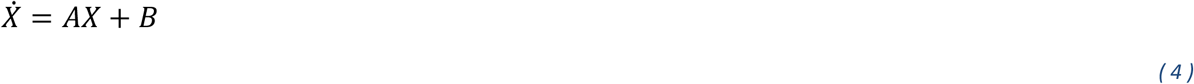

Thus, the ODE solver can’t calculate 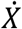 by matrix multiplications and may need to calculate each 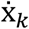 separately and save them into the matrix 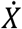 which increases computation cost exponentially by enlarging the size of the ODE system.

To overcome this challenge, the matrix of total reaction rates, denoted by *V* is calculated as

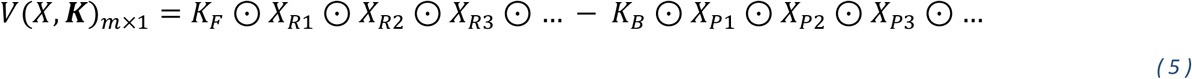

where *⊙* is the element-wise product and *K*_*F*_ and *K*_*B*_ include forward and backward reaction rate constants. *X*_*Pj*_ and *X*_*Rj*_ include the list of *j*_*th*_ products and reactants for all reactions respectively. The number of 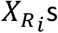 s and 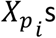 s depend on the maximum number of reactants and products of the reactions. By analyzing the roles of each state variable in reactions, matrix *H* is given by

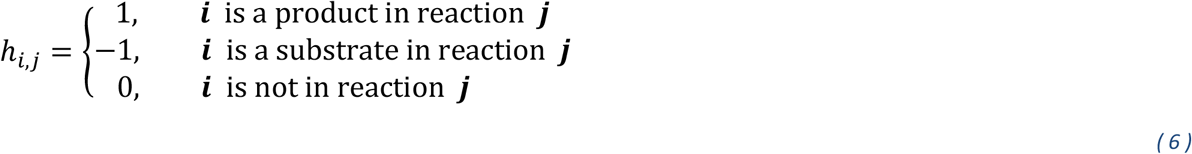

Consequently, multiplying *V* by *H* results in the matrix 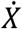 which is shown in (7).

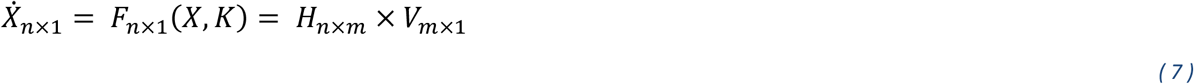

In this method of calculating 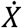, in each time step of integration, the ODE solver calculates indices of 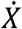 simultaneously. Testing this method on the nonlinear ODE model of colon cancerous cells with 3,347 state variables (which is explained further) shows that the proposed method could decrease mean calculation time from 2.5 seconds to 0.25 second for each time of ODE solving with similar initial values.

Taken together, using the SPADAN algorithm, the total biochemical reactions network is modeled by ordinary differential equations which have a set of unknown reactions’ kinetic parameters denoted as *K*. In further steps, SPADAN tries to find an acceptable approximation of *K* using time-course experimental data and parameter estimation techniques.

### Step IV: Parameter estimation

In the parametric ODE system automatically made by SPADAN, there is one parameter for one-directional and two parameters for two-directional biochemical reactions. Therefore, in large-scale models SPADAN works with, there is a large number of unknown parameters. Considering that Proteins, post-translational modified proteins, DNAs, and RNAs are the basic elements of the model, due to the shortage of experimental data on their biological reactions rates, providing acceptable estimations of model parameters based on time-series measurements is of utmost importance.

In order to estimate model parameter values, SPADAN uses the Least-squares method in which the gap between model simulation time points and experimental data time points is defined as an objective function called *S*. After that, model parameters are altered by an optimizer to minimize the objective function (REF). The way of calculating *S* is shown as follows:

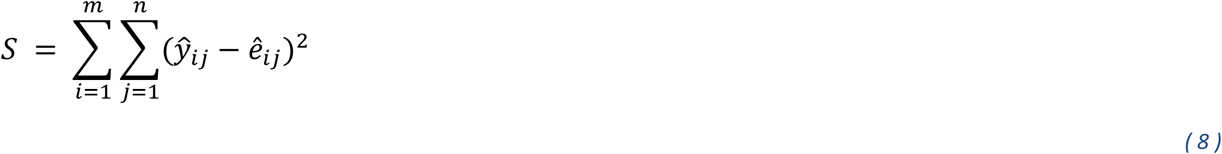

where ê_*ip*_ is the normalized experimental data and ŷ_*ip*_ is the normalized model simulation. The variables *i* and *j* show the number of biological elements and time points respectively. The total number of biological elements and time points are also shown with variables *m* and *n*. The normalization method of experimental data and model simulations depends on the measurement technique and the nature of measured values.

Due to the nonlinear kinetics of biochemical reactions, the ODE systems that SPADAN works with are mostly nonlinear. Therefore, during developing SPADAN, the performance of nonlinear optimization algorithms such as unconstrained quasi-newton, Nelder-Mead Simplex Method [11], and Levenberg-Marquardt Algorithm [12] was tested for large-scale parameter estimation problems. The results have shown that the progression toward the optimization solution becomes exponentially more complicated by increasing the number of unknown parameters and model equations. This is because gradient-based algorithms such as quasi-newton and Levenberg-Marquardt use numerical methods to estimate gradients which need numerous times of model simulations by increasing the number of unknown parameters. In addition, by increasing the size of the ODE system, each model’s simulation time increases which result in incrementing the total computation cost. However the Nelder-Mead Simplex Method doesn’t use derivatives, the number of needed model simulations for each step progression in it is related to the number of unknown parameters which makes it hard to find an answer in a feasible time.

To overcome this challenge, an approximation method has been proposed that speeds up the optimizer progression by considering the interconnectivity between parameters and state variables in ODE equations. Based on this method, the total optimization problem is broken into many sub-optimal problems to find the best possible answer. This method is the upgraded version of the optimization method declared in [13].

At the first step of this method, the parameters that exist in the derivative equation of each state variable are grouped into a set labeled with that state variable’s name. For example in the ODE system shown in Fig. 4, *K*_*xn*_ is the name of parameters related to the variable *x*_*n*_. In the next step, state variables are sorted based on their repetition time in the equations which is related to their effectiveness on the manner of the ODE system. In step 3, the Nelder-Mead optimizer starts to find the optimum S by altering the parameters belonging to the first group only. In the further steps, the optimizer performs this procedure for the next parameter groups. After reaching the last group of parameters, the algorithm starts from the first parameter group again. This procedure continues until getting to the proposed termination tolerance. The details of this optimization algorithm are shown in Fig. 4. In summary, the proposed approximation algorithm finds an approximation for parameter values of the model using time-course experimental data.

**Fig. 4.**
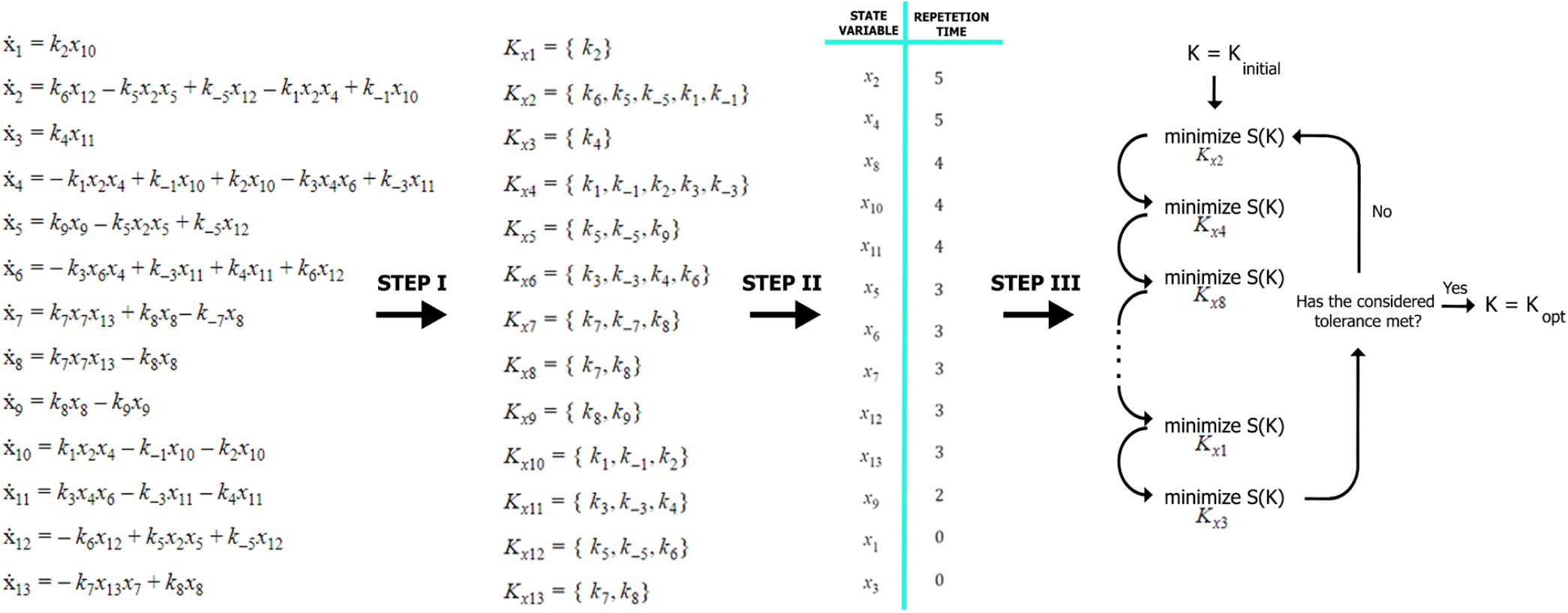
An example from the proposed parameter approximation algorithm. In the first step of the algorithm, one parameter group is made for each state variable. The groups named as 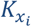 contain parameters exist in 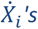’s equation. In step II, the algorithm sorts the state variables by the number of times each state variable is repeated in total state equations. During step III, the optimizer starts solving a sub-optimal problem from state variable with bigger repetition time to state variable with smaller repetition time. The sub-optimal problem for the state variable x_i_is to optimize the value of S in the space of parameters belonging to 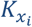. The algorithm is finished by reaching the considered tolerance of S.

### Case Study: Applying SPADAN to develop a dynamic model of colorectal cancer

In order to assess the applicability of the developed algorithm to construct a dynamic model based on a time-course experimental data, we have here re-analyzed and exploited a time-course multi-layer expression profiling data originally generated by Ressa et al [10]. These investigators assessed the response of WiDr colorectal cancer (CRC) cells to vemurafenib and gefitinib as BRAF inhibitor (BRAFi) and EGFR inhibitor (EGFRi), respectively. The cells were harvested 0, 2, 6, 24, and 48 hours after treatment and transcriptomics, proteomics, and phosphoproteomics data were generated. They found the up-regulation of metabolic pathways and tyrosine kinases receptors under BRAF inhibition as a primary response. Also, the switching of energy sources in treated cells turns to a defensive state to compensate for MAPK signaling inhibition. Noteworthy, they extended the analyses to a PTPN11 knockout WiDr cell line, the data of which is not used in the current study. The data of four experimental groups, including no treatment control, cells treated with BRAF inhibitor, EGFR inhibitor, or both are here explored.

Using MaxQuant analysis and after filtration, we identified 5655, 3432 proteins and phosphoproteins respectively. We were interested to consider the role and interactions of all identified genes and not only differentially expressed ones. Hence, using SPADAN, the map of interactions between all identified proteins, phosphoproteins, and transcripts in WiDr CRC cells was constructed (Fig. 5). Next, the biochemical reaction network was constructed which includes 5953 reactions and 3347 elements. The biochemical reaction network was then converted to a large ODE system with 3347equations and 7743 parameters.

**Fig. 5.**
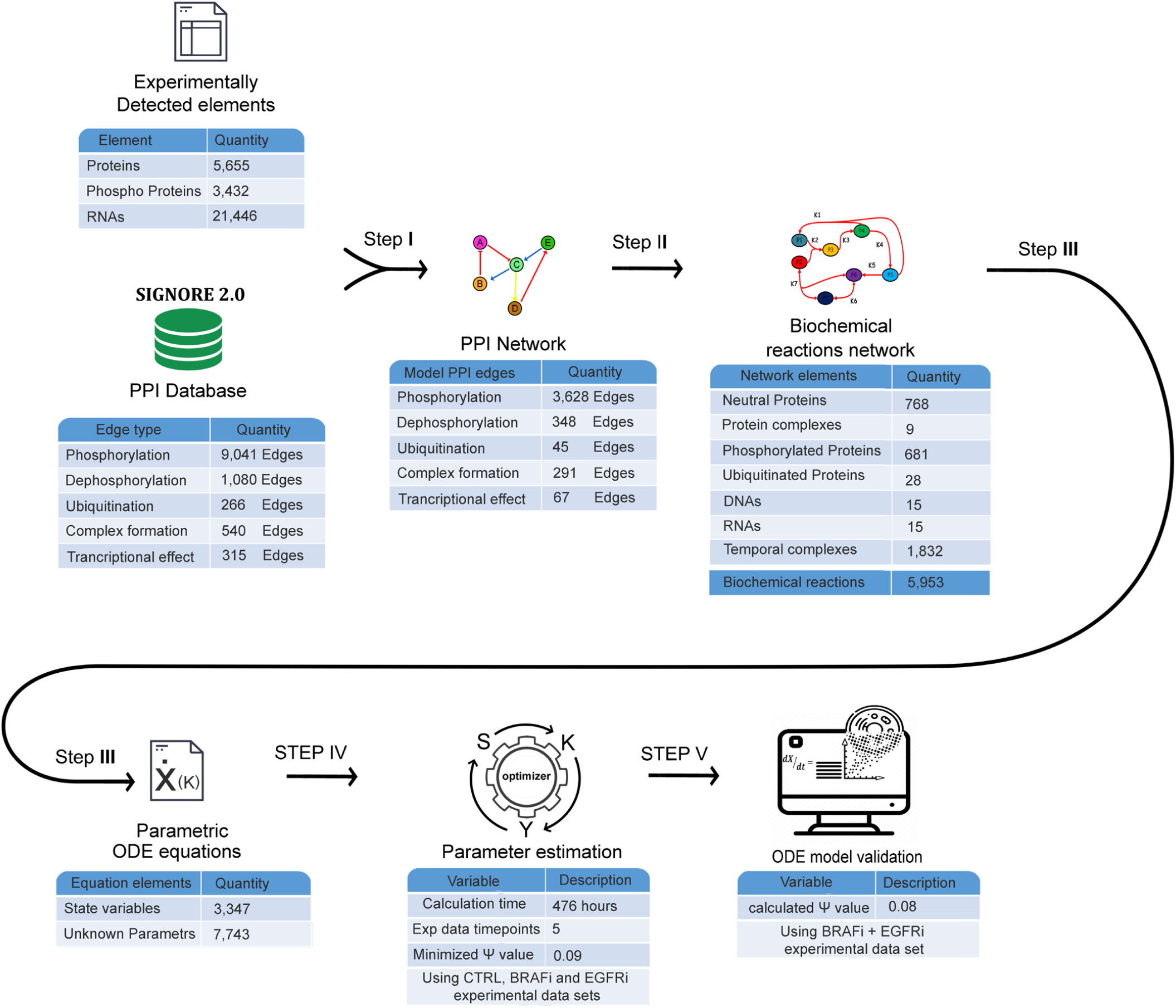
Statistics of case study modeling and validation. The experimental data includes 4 datasets called CTRL, EGFRi, BRAFi, and EGFRi+BRAFi. The first 3 datasets were used for modeling and the fourth was used for model validation.

#### Quantification of biomolecules

Proteomics data are generally expressed as relative quantifications using fold change parameters. However, the dynamic model simulation outputs are absolute concentrations. In order to make the comparison of these two feasible, the absolute concentration of proteins was estimated using the “proteome ruler” plugin which uses histones as standards. Employing this technique is not feasible for the estimation of absolute quantity of phosphoproteins. Hence, in order to compare phosphoproteomics experimental and simulation data, mean normalization was performed for both datasets; intensities of each phosphoprotein were scaled to make the mean value of the five time points equal to one. In order to calculate RNA absolute concentrations, normalized RNA counts were divided by an estimated volume of a cell (which is 2E-9 uL).

#### Approximation of basal concentrations

In ODE systems, initial values play an important role in the dynamic behavior of the system. Each protein concentration obtained from proteomics data is indeed the sum of concentrations of different states of that protein including phosphorylated form. In order to have an estimation of initial values of proteins and phosphoproteins, we relied on a previous study indicating that phosphoproteins constitute about 30% of total protein concentrations. Hence, 30% and 70% of the measured concentration of each protein at time zero was considered as the concentration of the phosphorylated and unmodified forms of that protein, respectively. We appreciate that in real situations this 30/70 ratio is not exactly true for all proteins. However, in the lack of absolute quantitative data, especially for phosphoproteins, this approximation can be acceptable.

For transcriptomics data, absolute concentrations are available for both time 0 and other time steps as indicated above. For the genes with transcriptional regulation in the model, DNA level is also included which can be bound to transcription factors or in the unbound state. The total DNA concentration obviously remains constant and is equal to 2 copies divided by cell volume which is about 8E-7 µL (REF).

### function S

In order to adjust model parameters, it is essential to compare model outputs with experimental data. In the exploited experimental data, measurements were performed with three biological replicates. The matrix of three biological replications for each time point of experimental data is denoted by 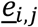 as follows:

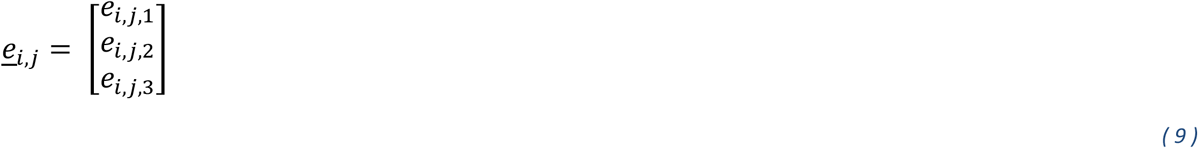

Where *i*is the number of gene, *j* is the number of timepoint and the third index represents the number of biological replicates.

Since in this case study, the experimental data is from proteins, phosphoproteins, and mRNAs, the concentration of proteins, phosphoproteins, and mRNAs of each gene in the model are defined as model outputs. As mentioned before, each molecule can appear in different biochemical elements. Therefore, in order to find the intensity of a molecule at a time point, the concentration of these elements should be summed. *Y*_*P*_, *Y*_*Phospho*_ and *Y*_*RNA*_ are three sets of model outputs which show the value of proteins, phosphoproteins, and mRNAs in the model respectively. These values are calculated by adding their element values which are calculated by multiplication of the C matrix to the X. C matrix consists of three binary matrices called *C*_*P*_, *C*_*Ph*_ and *C*_*RNA*_ which are automatically made by SPADAN.

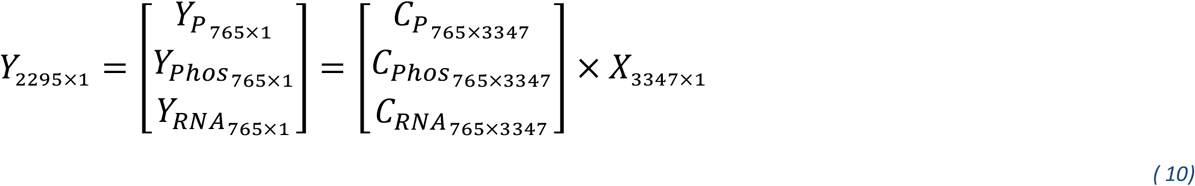

Considering the wide range of concentration scales, the sum of the squares of differences between model output and experimental data in different time points is calculated for each element and then divided by the average of experimental concentrations in the five timepoints.

For proteomic and transcriptomic experimental data which there were estimations for their concentrations, the average of experimental concentrations in the five timepoints 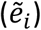 is calculated by:

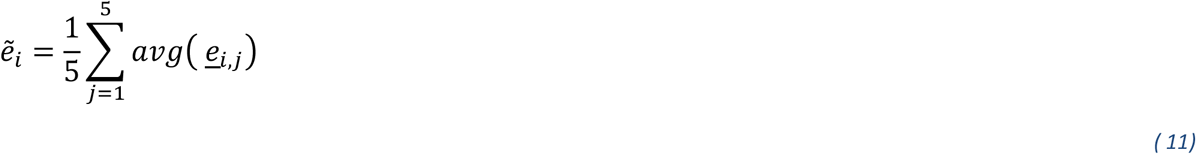

For each time point, if the simulation value (*y*_*i,j*_) is in the range of minimum and maximum of three experimental replicates which are shown with *max(*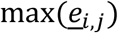) and *min(*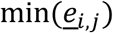 *), a*_*i,j*_ gets zero and the difference between simulation and experimental data is ignored in calculating S. Otherwise, the residual between simulation data and mean value of the three replicates (*d*_*i,j*_) was used to calculate function S as the sum of squares of differences between simulation and experimental data.

Variable *a*_*i,j*_ is calculated using:

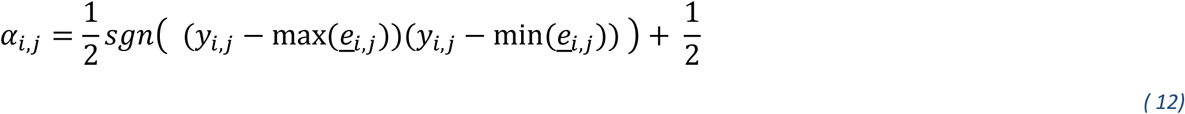

To calculate S relatively, it is calculated as follows:

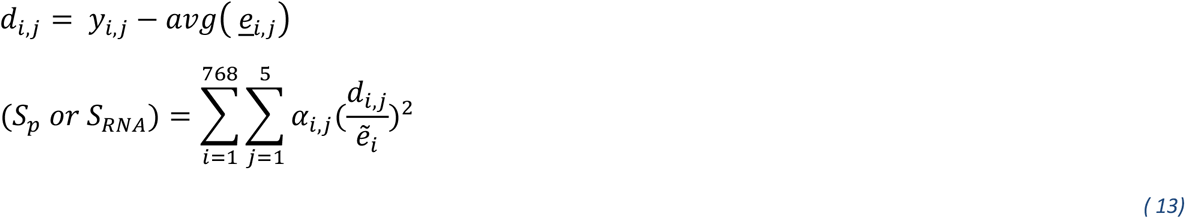

where *S*_*p*_ and *S*_*RNA*_ are the sum of squares in proteomics and transcriptomics levels respectively, *i*is the gene number and *j* is the number of timepoint. A visual example of this normalization and S calculation method is shown in Fig. 6.

**Fig. 6.**
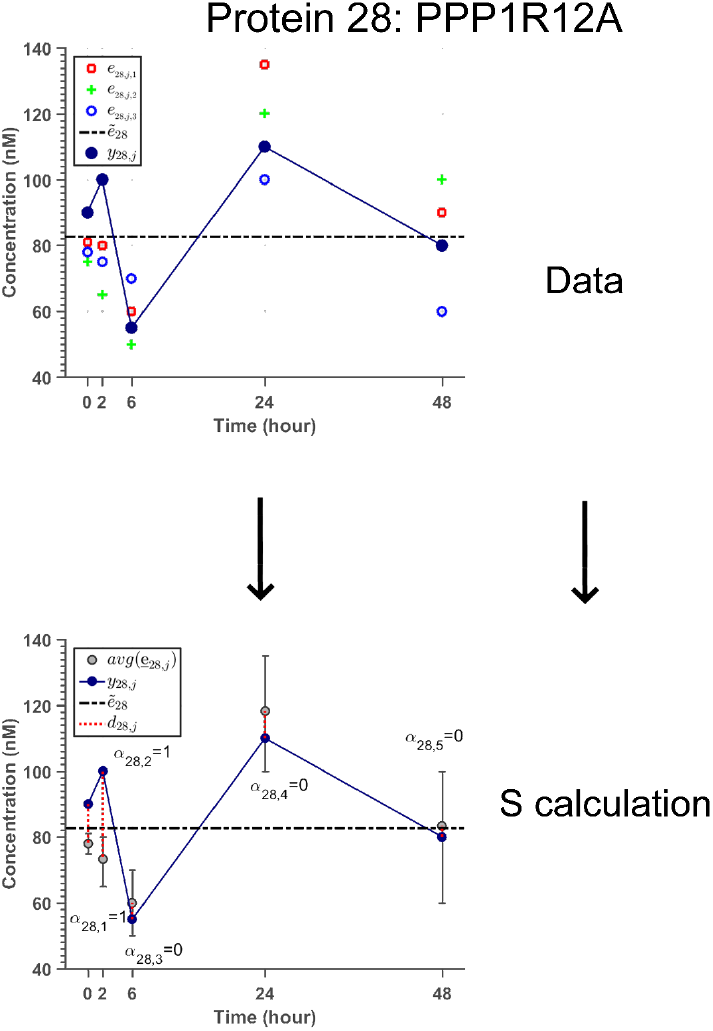
An example of comparing method between model simulation and experimental data for RNAs and proteins. Three biological replications for each experimental data time point are shown with red, green, and blue markers. Since both experimental data and model simulation are absolute concentrations, there is no need for mean normalization. The method of calculating S is explained in (11) to (13).

Since in this case study, there was not a method to estimate the absolute concentration of phosphoproteomics experimental data, mean normalized intensities and model simulations were used to compare the track of concentration changes during the time. Mean normalization for experimental intensities are calculated as follows:

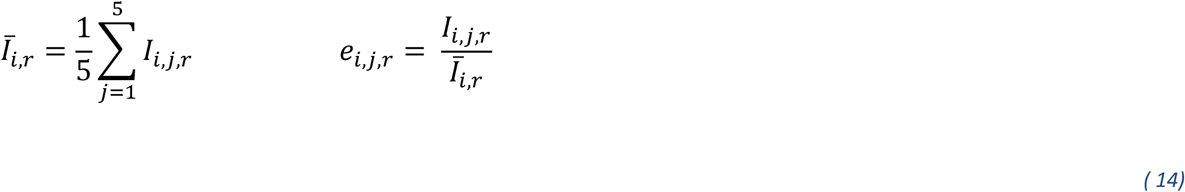

*I*_*i,j,r*_ is the intensity of the *i*_*th*_ phosphoprotein at the *j*_*th*_ timepoint in the *r*_*th*_ biological replication. After defining *e*_*i,j,r*_ in (14), the matrix 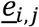 is constructed as (9).

Model simulations are also mean normalized by:

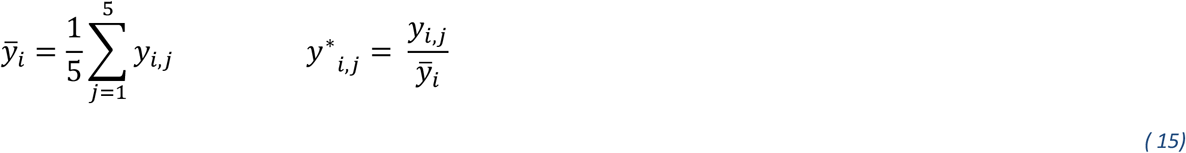

where *y*^*∗*^_*i,j*_ is the mean normalized simulation value of the *i*_*th*_ phosphoprotein at the *j*_*th*_ timepoint.

In the following, the sum of squares in phosphoproteomics level which is shown with *S*_*ph*_is calculated by:

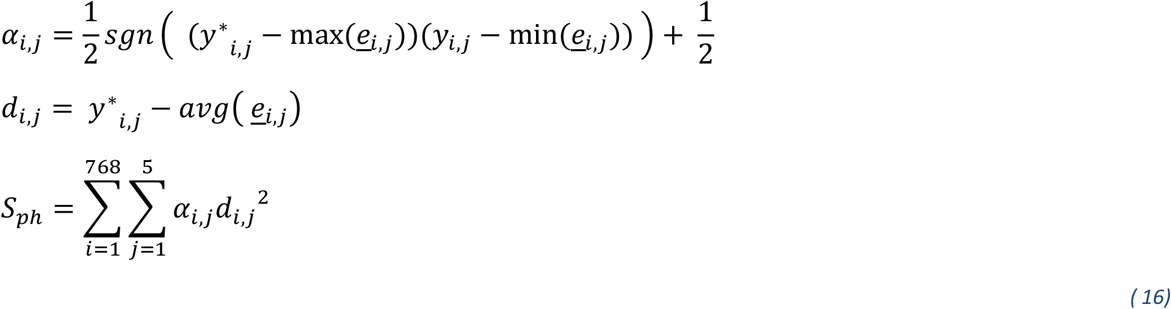

in which *d*_*i,j*_ is ignored in the summation if *α*_*i,j*_ is zero.

Obviously, this normalization applied for proteomics and transcriptomics data is not required for phosphoproteomics data which were previously mean normalized. The normalized S for all model elements are then summed to calculate *S*_*total*_ in the equation below:

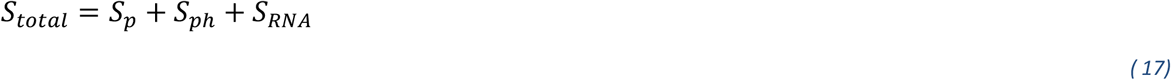

A visual example of S calculation in phosphoproteomics level is shown in Fig. 7.

**Fig. 7.**
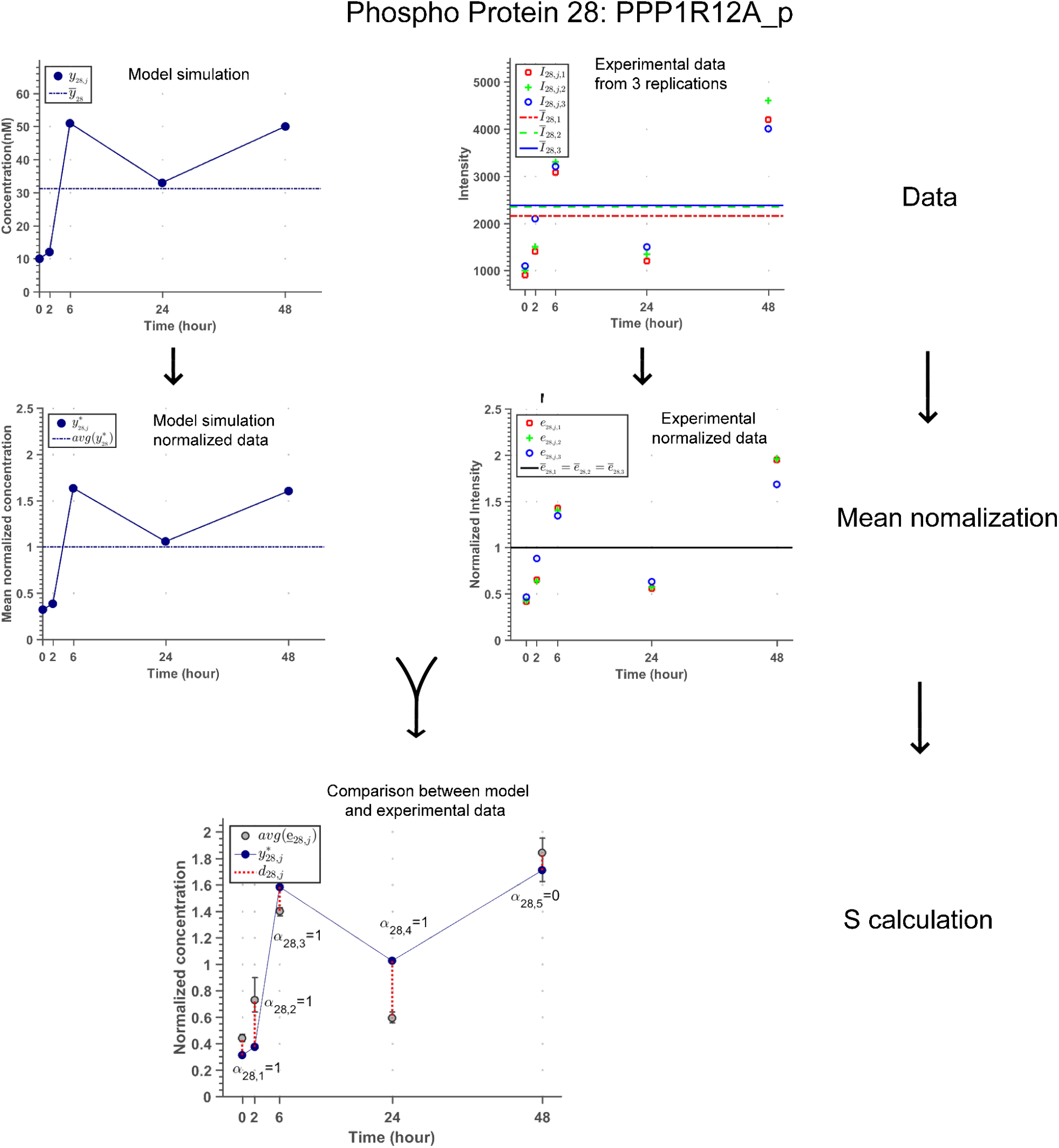
An example of comparing method between model simulation and experimental data for phosphoproteins. Since experimental data from phosphoproteins are intensities and model simulations are concentrations, they are both mean normalized to get comparable. Each of the biological replications is shown with red, green, and blue markers. The method of calculating S is explained in (14) to (16).

Using the SPADAN parameter approximation algorithm to approximate unknown model parameters, could decrease *S*_*exp*_ from *2*.*72 × 10*^*7*^to *1*.*13 × 10*^*6*^ after 476 hours of calculation. In order to get a better nsight about the mean value of S for each timepoint of the model, *Ψ* is declared in (18) which is the mean value of Root Summed Squared (RSS) of the model residual between *5* timepoints of *768 × 3* biomolecules. The progression of the proposed optimizer in minimizing *Ψ* from initial guess to approximated parameter values is shown in Fig. 8.

**Fig. 8.**
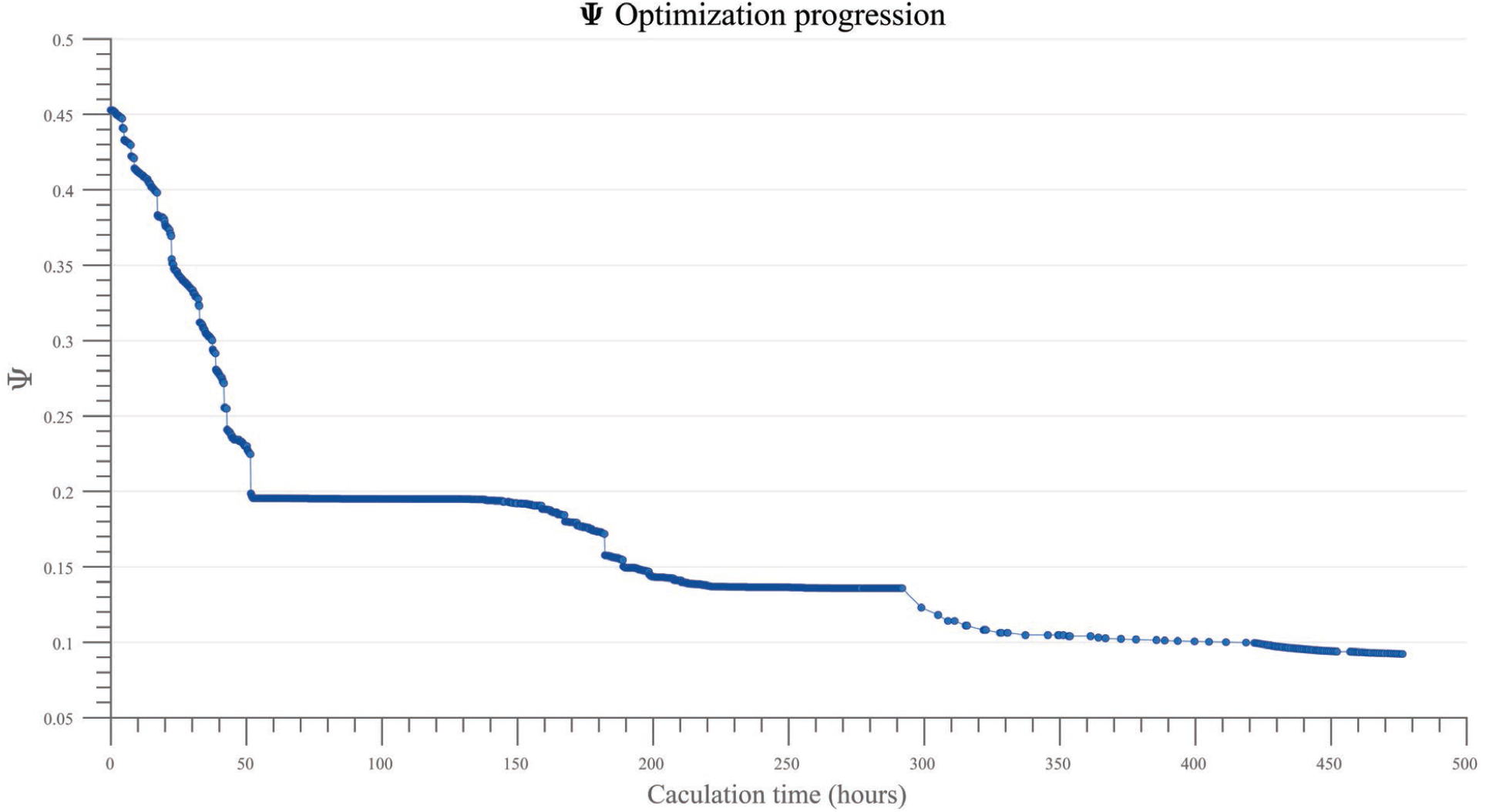
Optimization progression during the time of calculation. By applying the proposed parameter approximation method, after 476 hours of calculation, the S value was decreased from 2.72 × 10^7^to 1.13 × 10^6^ which equals decreasing Ψ value from 0.45 to 0.09 for each time point.

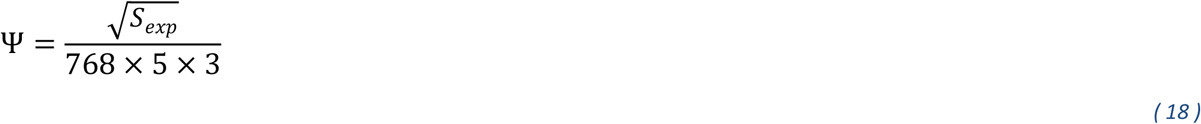

### Model validation

After estimating model parameters to fit outputs to experimental data from control, EGFRi, and BRAFi groups, the model was run in the situation of simultaneous treatment with EGFRi and BRAFi drugs. Simulation results were then compared with experimental data to assess the accuracy of the model prediction. The *Ψ* value which shows the difference between model prediction and the experimental data for BRAFi + EGFRi dataset was *0*.*083* indicating that the model could present an acceptable prediction. The comparison of model simulations and experimental data for some representative biomolecules is shown in Fig. 9. The figure shows that for genes such as NFKB1 and SF1, the model could predict the trajectory of concentration changes during the time, considering the model’s weakness in predicting concentration changes for genes like SLC2A1 and GDF15.

**Fig. 9.**
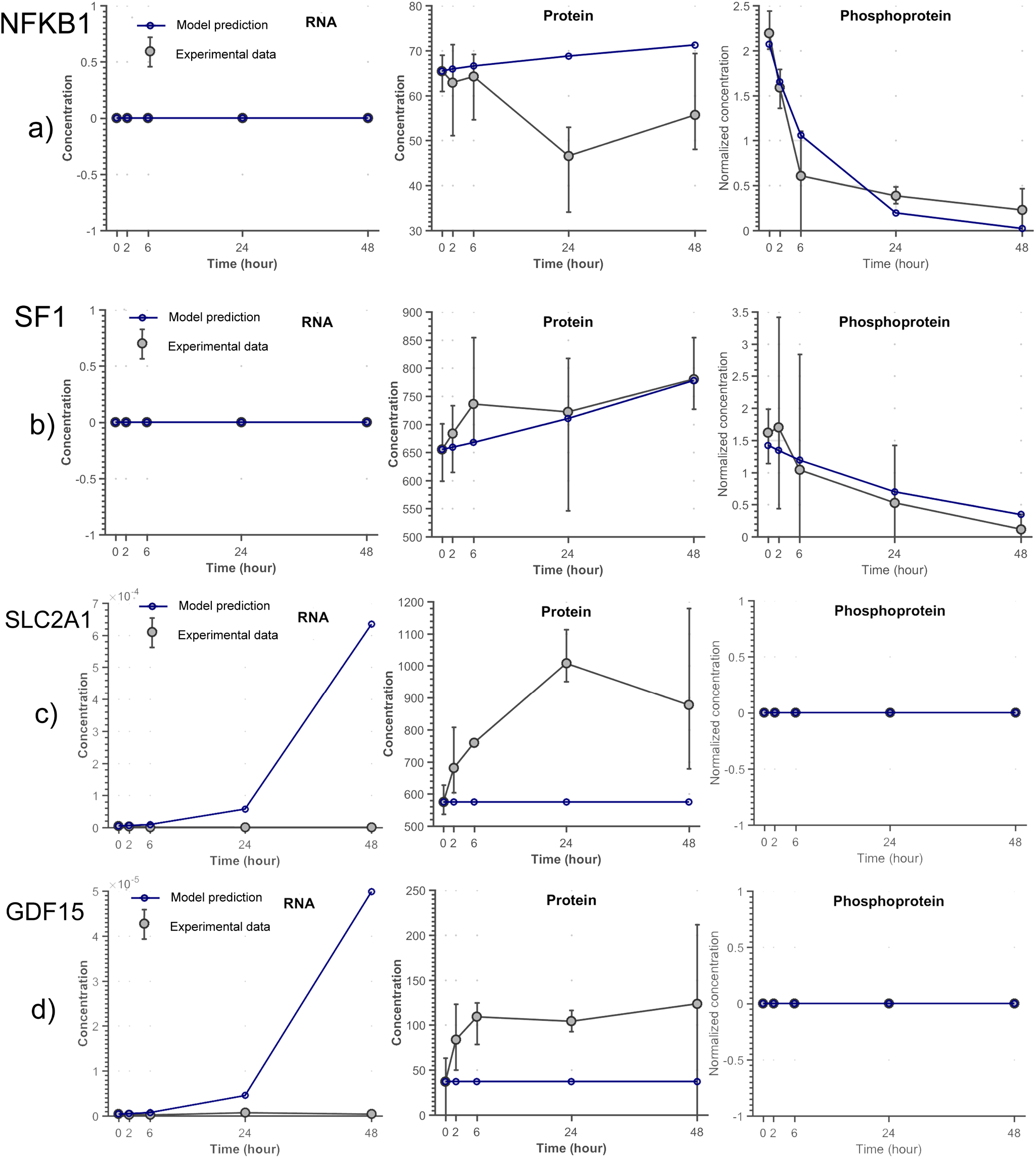
Model prediction vs. experimental data for some representative genes from response of WiDr colorectal cancer (CRC) cells to simultaneous Vemurafenib and Gefitinib treatment. Gray circles show the mean value and error bars show the minimum and maximum values between three biological replications. However the model could have an acceptable prediction from the manner of genes such as NFKB1 and SF1 shown in figures a and b, for genes such as SLC2A1 and GDF15 it could not have a precise prediction (figures c,d).

## Discussion

Despite invaluable insights provided by systems biology in the last few decades, a major unmet flaw is that the constructed models are either holistic or dynamic (REF). In the top-down approach, big data are organized to generate holistic but static maps of the interactions. On the other hand, in the bottom-up approach, mathematical predictive models can be constructed that incorporate the cell dynamism but ignore many critical elements and focus on limited numbers of role players. An appropriate response to the insufficiency of current therapies for most non-communicable diseases requires large-scale dynamic models. The current study was aimed at the development of such a framework.

A bottleneck in construction of big dynamic networks is to identify biochemical reactions involved in the interaction between two biomolecules. Indeed, every single edge between two proteins in a PPI network is a compact code that should be decoded to the more comprehensive language of biochemical reactions. We have generated a conversion list that could be assumed as a “dictionary” for the translation of graph edges to reactions. Biochemical reactions are then converted to a set of ODEs. Hence, state-space equations can be extracted from large-scale networks using the above steps. Although in the presented case study, all equations are based on mass action, SPADAN allows to employ Michaelis-Menten or Hill kinetics as well.

In this study, we have employed a time-course large scale experimental data on colon cancer cells to assess the validity of the developed framework. An advantage of the constructed network is that it encompasses different layers of biomolecules. Specially, it includes all identified proteins which is in contrast with most studies that focus solely on the interactions of differentially expressed proteins. This common strategy can be scrutinized by ignoring the interactions of proteins that are not differential but can be involved in critical interactions.

Constraint-based modeling allows the generation of a “solution space” for the rates of biochemical reactions that are shrunk by imposing a variety of constraints. Developed in the mid-twentieth century, this strategy has been more recently employed for modeling large-scale biochemical systems [7], [14], [15]. However, it cannot provide detailed insights about the concentrations of biomolecules and the exact kinetics of each reaction. The construction of large scale dynamic models has been attempted by a few previous investigators; Smallbone et al have developed a strategy for the construction of genome-scale dynamic models of metabolic networks and employed it to generate a holistic kinetic model of yeast metabolism. In spite of the high merit of this work, the parameters were harvested from a kinetic model repository. This makes its application limited to cases that such a library is already available. Similarly, Smith et al have developed a computational framework that receives an SBML format network and automatically constructs differential equations for biochemical reactions. The reaction rates are harvested from several databases including previous measured parameters. Notably, these investigators have shown that accurate estimates for more than 80% of reaction rates are required for an acceptable simulation. This is too far from what is already available[5]. This indicates that in the deficiency of such experimental knowledge, parameter estimation is at the cornerstone of large-scale dynamic modeling of biological systems.

The parameter estimation step can be assumed as the major limitation hindering the construction of large-scale ODE models. This procedure is performed by minimizing the differences between trajectories of model outputs and time-course experimental measurements [16]. Although a variety of optimizers are available for this purpose [17], they could hardly be applied for large-scale ODE models. Therefore, in this study, a novel parameter approximation tool is developed and incorporated into SPADAN which fits with the complexity of equations and the large dimensions of genome-scale models.

Considering the complexity and nonlinearity of biological systems and the scarcity of time-course experimental data, the parameter estimation strategies, including the method employed in this study, may result in a set of parameters that is not unique. We propose that integrating the core concepts of constraint-based modeling with the introduced parameter estimation strategy may limit the space of parameters and hence result in more accurate approximations. This remains to be examined in future works. Additionally, to fasten the parameter estimation speed, the proposed parameter estimation algorithm has the potential to be parallelized which makes it computable on multi-core supercomputers. This enhanced computation approach would pave the way for more sophisticated analyses such as considering the stochasticity of biological reactions. Additionally, the introduced large-scale modeling can be improved by considering the intracellular compartments/spaces that the reactions take place as well as intracellular transports.

In conclusion, we have proposed a modeling and optimization method which can fill the gap between large-scale static and small-scale dynamic modeling strategies. This simulation scheme allows quantitative analysis of cell behavior and prediction of response to different therapeutic interventions which is a major step towards precision medicine.

## Footnotes

1 http://proteomecentral.proteomexchange.org

